# Poor decision-makers: motivation, working memory performance, and repartition across two inbred strains of rats

**DOI:** 10.1101/2022.06.07.495184

**Authors:** Aurelie Y. Fitoussi

## Abstract

A minority of healthy individuals (poor decision-makers, PD) exhibit a combination of behavioral traits reminiscent, at least in part, of addiction and predicting poor decision-making (DM), namely motor impulsivity, inflexibility, risk-taking, and higher motivation in Wistar Han rats. Two behavioral features, motivation and working memory (WM), play a role in DM capacities although the precise relationship is not entirely known. Additionally, we previously reported that neurotransmitters e.g., dopamine - modulation was tightly linked to the PD phenotype. The goal of the study was to investigate the detailed motivational functions in PD individuals including saccharin intake, reward-seeking or incentive behaviors under different internal states i.e., food-deprived or *ad libitum*. Maze-based spatial WM was also evaluated. Moreover, two inbred strains of rats, Lewis and Fisher 344 (F344) rats, known for modeling vulnerability to drug addiction and affected by substantial variations in the mesolimbic dopaminergic pathway, were run in the DM task (Rat Gambling Task, RGT). PD Wistar Han rats displayed higher saccharin intake levels and a drastic increased reward-seeking behavior on a fixed schedule. PD were more sensitive to the internal state in responding to saccharin delivery in fixed but not in progressive schedules. A few relationships were found within motivational functions, and with DM, that is a positive correlation between saccharin intake and reward-seeking behavior, and a negative correlation between saccharin intake and DM. PD were significantly not impaired in WM. Lewis and F344 rats displayed improved performance early in the task (exploration) and a higher proportion of PD was observed in Lewis as compared to F344 rats. Altogether, these findings complete the preclinical panel of behavioral functions that relate to poor DM and extend a presumed role of dopamine in such processes.

## Introduction

Real-life decision-making (DM) is impaired in several psychiatric conditions but also in a subset of healthy individuals for whom high immediate gratifications prevail over long-term gain in humans and rodents (Rivalan et al., 2009; Bechara et al., 1994). Psychological features and cognitive processes related to DM capacities have been studied in pathological states (Kovacs et al., 2017; Brevers et al., 2014) while healthy individuals with poor DM performance are largely neglected (Suhr and Tsanadis, 2010; Suhr and Hammers, 2006). DM results from the integration of executive functions that allow for solving complex tasks. Thus, higher-order cognitive processes including planification, inhibition, and deduction abilities are essential components to achieving good decisions in complex and conflictual situations (Chudasama et al., 2006; Royall et al., 2002). It is noteworthy that there is no consensus yet as to what extent working memory performance relates to DM (Brand et al., 2006). Obvious pieces of evidence however are in favor of a non-direct relationship between these cognitive domains, and clinical studies indicated a possible functional link between working memory and DM, especially in addict patients (Bechara and Martin, 2004).

A Rat Gambling Task that tracks DM capacities like in the above situation, closely mimics the same principle of the Iowa Gambling Task, a human model of real-life DM (Rivalan et al., 2011; Rivalan et al., 2009; Bechara et al., 1997). Like in humans, a minority of healthy Wistar Han rats persist to choose the disadvantageous options. Theoretically, it has been hypothesized that a continuum between the normal and the pathological state emerges along with possible behavioral dimensions such as the impulsivity and compulsion dimension (Dellu-Hagedorn et al., 2018), common aspects relevant to addiction (Koob and Volkow, 2010). In this framework, categorized maladapted - poor performers as “healthy” individuals could share certain behavioral and neurobiological features as pathological states, raising eventual endophenotype identification and/or vulnerability markers to DM-related disorders (Gottesman and Gould, 2003). It has been shown that these poor aforementioned decision-maker individuals exhibit a combination of behavioral traits reminiscent of addiction, namely motor impulsivity, inflexibility, risk-taking, and higher motivation in some circumstances that are good predictors of DM performance (Rivalan et al. 2013; Rivalan et al., 2009). These poor decision-makers (PD) also displayed impaired goal-directed behavior (Fitoussi et al., 2018), a characteristic tightly coupled with the development of drug abuse (Vandaele et al., 2018), but preliminary data favored no enhanced drug responsiveness in self-administration models. At the neurobiological level, PD recruit a limited prefrontal-subcortical network while performing the RGT (Fitoussi et al., 2015), with some common identified neurobiological substrates relevant to executive functioning, such as the orbitofrontal cortex and medial prefrontal cortex (Costa et al., 2021; Keiflin et al., 2013; de Visser et al., 2011; Balleine and O’Doherty, 2010). It is important that key subcortical brain regions including the nucleus accumbens (core) and the amygdala (basolateral nucleus) play a critical role in such behavioral phenotypes pointed out a major involvement of motivational functions (Zeeb et al., 2013; Balleine et al., 2005; Berridge, 2004). It was previously observed that PD exhibit higher motivation after the manipulation of the reward magnitude during a progressive ratio schedule (Rivalan et al., 2009). Thus, additional features underlying this phenomenon may be responsible such as metabolic and cost variables (Baldo et al., 2013).

Aside from a limited network sustaining poor DM capacities during the RGT, neurotransmitter levels have been shown to relate to poor decisions, especially dopamine action in cortical and subcortical brain regions including the infralimbic cortex (a subpart of the medial prefrontal cortex) and nucleus accumbens (Fitoussi et al., 2015; Cardinal et al., 2001). However, a causal relationship has not been investigated. In this respect, two inbred strains of rats are of particular interest, namely the Lewis and Fisher 344 (F344) rats. Indeed, the Lewis rats are a good candidate for their ability to sustain self-administration of drugs of abuse (Cadoni, 2016), a behavioral feature that has been linked to dopamine level modulation. Additional characteristics of the dopaminergic system have been reported including those related to dopamine metabolism and neural activity (Haile et al., 2001; Harris and Nestler, 1996; Minabe et al., 1995). Anxiety level and stress differences could also be described in these rat models (Ramos et al., 1997; Chaouloff et al., 1995).

The goal of the present study was twofold: 1) to investigate the motivational functions of PD measured in the RGT including saccharin intake in a free-choice procedure, reward-seeking and incentive behaviors in fixed ratio 5 (FR5), and progressive ratio (PR) under food-deprived or *ad libitum* condition, as well as the eventual relationship between working memory measured in an 8-radial maze and DM capacities; 2) the repartition of PD among two inbred strains of rats, Lewis, and F344 rats.

## Material and Methods

### Animals

Male Wistar Han, Lewis and Fisher (F344) rats (Charles River, Lyon, France) aged from 13 to 15 weeks were used. They were housed in groups of four in a temperature-controlled room (22 °C) on an inverted 12-h light/dark cycle (light on at 08:00 p.m.). Tests were conducted during the dark phase of the cycle. They had free access to water and were moderately food-deprived (95 % of free-feeding weight) when required throughout the experiments. All procedures were conducted in strict accordance with the 2010–63-EU and with the approval of the Bordeaux University Animal Care and Use Committee (Permit Number: 5012087-A).

### The RGT. Behavioral apparatus and procedures

Twelve identical operant chambers (Imetronic, Pessac, France; adapted from 5-choice serial reaction time chambers) were used for the Rat Gambling Task (RGT). Four circular holes were available on a curved wall and could be dimly illuminated with a white light-emitting diode (LED) located at their rear. A food magazine on the opposite wall was connected to an external dispenser delivering food pellets (45 mg, formula P, TestDiet, USA). A clear vertical Plexiglas partition (Imetronic, Pessac, France) (28 cm × 9.5 cm × 9.30 cm) with a central opening (7 cm × 9.7 cm) was placed across the middle of the chamber, parallel to the food wall. Data collection was automated using control software (Imetronic, Pessac, France) running on a computer outside the testing room. *Training*. Procedures were run as previously described (Fitoussi et al., 2015). During training (5–7 days), rats (n = 48) learned to associate a nose-poke visit with the obtention of a food pellet until 100 pellets were reached. If not, the session ended within 30 minutes. The next step was to associate 2 nose-poke visits with the obtention of a food pellet, the same criterion as above. The latter was defined to ensure that the visit of the animal was voluntary. Finally, animals were submitted to 2 additional sessions: (1) 2 nose-poke visits led to 2 food pellets delivered as the next RGT test session. A criterion of 100 food pellets or 15 minutes was defined, and (2) 2 nose-poke visits led to 1 food pellet as the next RGT test session. A criterion of 50 pellets or 15 minutes was defined.

### RGT test

After training, rats were tested in a 60-min RGT test session during which they could freely choose between 4 options (A–D) (exploration) before establishing their preference (exploitation). Choices C and D vs. A and B led to the immediate delivery of 1 vs. 2 pellets respectively, but choices A and B were disadvantageous in the long term since they could be followed by much longer, unpredictable penalties. Good decision-makers (GD) and PD were differentiated based on the percentage of advantageous choices (70 and 30 %, respectively) during the last 20 min of the test. Individual differences among GD were also analyzed. GD with a fast time course to make good choices (FAST GD), on average superior to 50% within the first 10–20 minutes, were counted as well as GD with a slower time course to make good choices (SLOW GD), on average inferior to 50% within the first 10–20 minutes. Undecided rats with intermediate scores were discarded because of the small number of rats in this category.

### Working memory, 8-radial maze. Habituation and training

On the first day, rats could freely explore each arm separately as well as the central platform for 1 minute. On the following day, rats were placed on the central platform with closed arms for 10 seconds. At the end of the delay, rats could freely explore the maze and each baited arm. Each time the rat returned to the central platform, the arm was closed. The session ended when all baited arms were visited. This was performed for 2 consecutive sessions.

### Acquisition

This step lasts 8 days, a session a day. Like the previous phase, each arm was baited, and the rat was placed on the central platform for 10 seconds before the arms were opened. The difference with the training was the fact that if the rat returned on a previously visited arm, all the doors of the maze were closed for 6s, and then re-opened. The session ended when all arms were visited or 16 visits counted (20 minutes, if not).

### Test, delay effect

During this step, sessions were similar to previously, but the delay between choices varied: 1s, 6s (like the acquisition), and 12s, a session per condition and per day. The criterion for ending the session was similar to the above.

### Saccharin intake, dose-curve effect

We ran a novel procedure. The procedure consists of 2 hours-daily sessions. 2 water bottles were placed on the right versus left side during the first session i.e., habituation. Then, the amount of liquid (water) consumed was monitored for 2 subsequent days as well as a spatial preference (a session a day). After validating an equal amount of consumed water, rats were then allowed to freely drink the water or saccharin solution (replacing one previous water bottle) at the following doses: 0.009% (0.5 mM), 0.018% (1 mM), 0.055% (3 mM), 0.09% (5 mM), 0.18% (10 mM), 0.36% (20 mM), 0.54% (30 mM), and 0.9% (50 mM). 2 consecutive sessions with stable consummatory behavior were required before switching to another dose together with bottle location (i.e., left or right side). The procedure ended with 2 sessions of water consumption only.

### Fixed-ratio 5, progressive ratio 2

The comparison of operant behavior and motivated-related behaviors was assessed using a FR5 and a progressive ratio 2 (PR2) procedures. During FR5, 5 lever presses were required to earn a dose of saccharin. The session ended within 30 minutes. During PR2, lever presses escalated throughout session 2 by 2 lever presses (ex: 2, 4, 8, 16, 32, etc.) to earn a dose of saccharin. The session ended within 30 minutes. In these procedures, the state need of the animal was assessed by either food-deprived them (same as before) or letting them *ad libitum*. The minimal dose of 0.055% of saccharin or the maximal dose of 0.18% was tested in each condition. 5 sessions per condition were required.

### General data analysis

Comparisons of behavioral scores (mean ± SEM) with random choice (50 %) in the RGT were made using a two-tailed t-test for groups and subgroups of GD and PD. Statistical analyses were then made using ANOVAs, e.g., comparison of scores time-course between groups (repeated measures, group and time factors), number of choices, number of visits, saccharin scores, and working memory parameters, followed by the Fisher’s PLSD post-hoc test when required (Statistica, Statsoft 7.0). Linear regression analyses were made using Pearson correlation (r).

## Results

### Inter-individual differences in decision-making

Distinct patterns of choice preference within a single RGT session were reported (Fig. 1). All rats (n = 19) sampled the 4 available options (DM, exploration) before establishing a preference (DM, exploitation) for A and B, or C and D options. They clustered into 2 distinct categories depending on their final preference: (1) a majority of GD (n = 13) with a strong preference for advantageous options (70 % preference); and (2) a minority of PD (n = 6) that persevered in choosing disadvantageous options (70 % preference). Importantly, and despite differences during the exploration phase of DM, all rats sampled the 4 options during the first 10 min and experienced the long penalties at least once. Among GD, rats could be distinguished according to the time course of DM. Some rats (SLOW GD, n = 6) still chose the options randomly during the first 20 min (comparison with chance level (50 %), t Student = 1.52, ns), whereas others (FAST GD, n = 7) promptly orientated their choices toward advantageous choices (Fig. 1A). The total number of visits (Fig. 1B) of GD and PD was similar (F_2,19_ = 2.77, ns). Consequently, the number of choices in either A, B, C, and D holes varied greatly between GD and PD. PD performed more A and B options than C and D options and related to GD, both FAST GD and SLOW GD (F_2,19_ = 23.75, p < 0.01) (Fig. 1C). Additionally, the pellet consumption of GD was largely superior to PD (F_2, 19_ = 17.89, p < 0.01) (data not shown).

**Figure 1:**
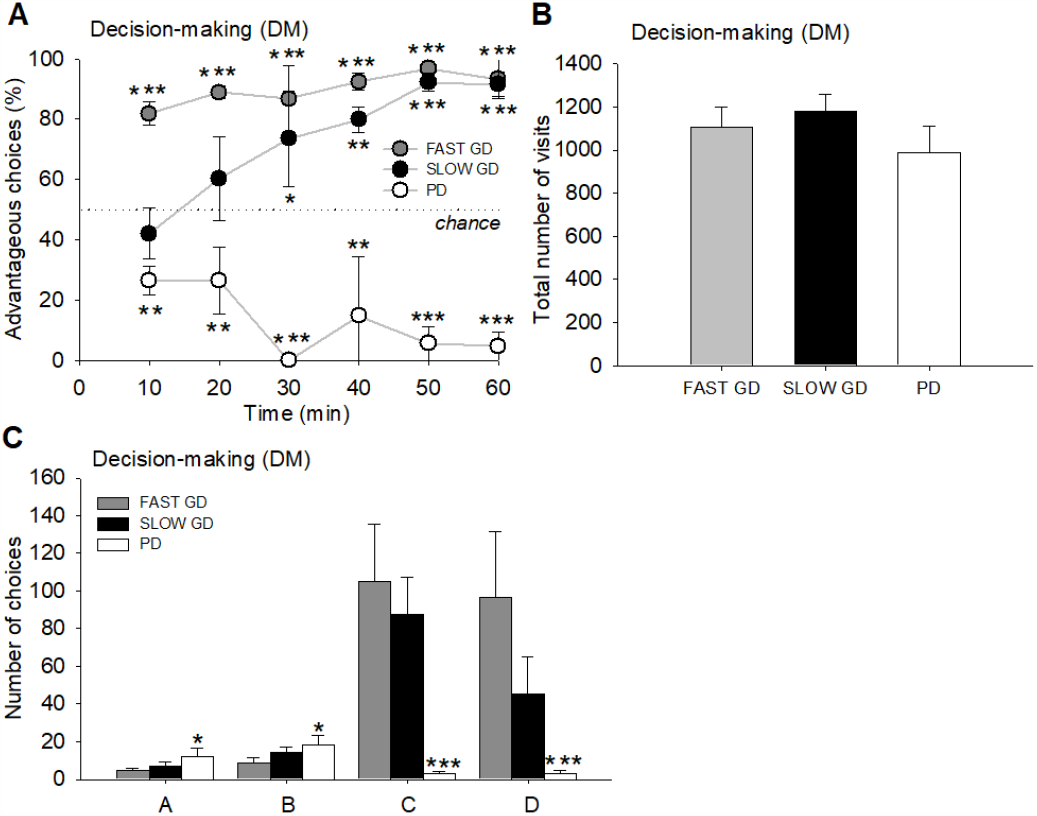
RGT performance. (A) Pattern of choices of good decision-makers (GD), FAST versus SLOW GD, and poor decision-makers (PD). SLOW GD displayed a longer time course to choose the advantageous options (especially during the first 20 minutes) than FAST GD. PD persisted to choose the disadvantageous options. (B) Total number of visits, and (C) Number of choices in the holes A, B, C, and D for the FAST and SLOW GD, and PD. GD preferred C and D choices. *\* p < 0*.*05, ** p < 0*.*01, *** p < 0*.*001, t-test Student or ANOVA when required*.

### Inter-individual differences in working memory

Inter-individual differences in the acquisition rule and working memory were assessed during the acquisition and delay phase (test) of the procedure. During acquisition (Fig. 2A, C, E), no difference in performance as shown by the total number of choices was revealed (F_2,19_ = 3.42, ns), as well as the percentage of errors/8 choices parameter (F_2,19_ = 4.90, ns). However, striking differences in the number of perseverations (returns to a previously visited arm) during this phase on day 4, day 6, and day 9 (F_2, 19_ = 31.41, p < 0.05; PLSD Fisher, p < 0.01) were evidenced. PD perseverated more than GD. During the delay phase (Fig. 2B, D, F), no difference in working memory performance as shown by the total number of choices (F_2,19_ = 6.75, ns) and the percentage of error/8 choices parameters (F_2, 19_ = 5.65, ns) was revealed, even if a not statistically significant trend was noted for the 12s delay duration. Finally, no difference in perseverations during the delay phase was reported (F_2, 19_ = 7.78, ns).

**Figure 2:**
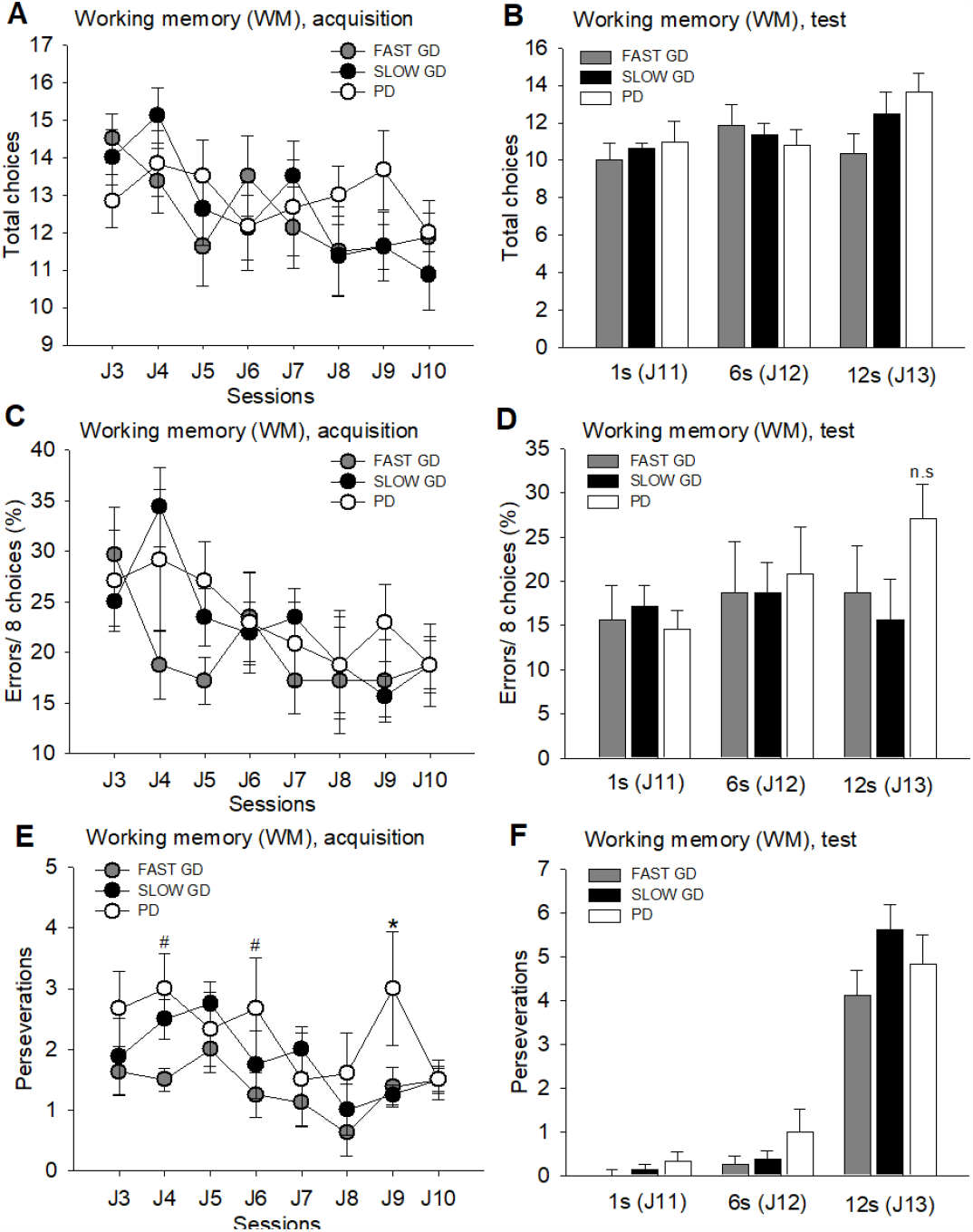
8-radial maze performance. (A) Total number of choices during acquisition in the working memory (WM) task, (B) Total number of choices during the test of WM during 1s, 6s, and 12s retention delay. (C) Number of errors for the 8 first choices during acquisition and (D) performance parameter i.e., number of errors for the 8 first choices during retention delays for FAST and SLOW GD, and PD. There was no difference between all groups. Additional parameters i.e., (E) number of perseverations during acquisition, and (F) during the test phase during retention delays for FAST and SLOW GD, and PD. *\* p < 0*.*05, # p < 0*.*05 (comparison between FAST GD and PD), ANOVA*.

### Inter-individual differences in saccharin intake, dose-curve analysis

During this phase, the basal consumption of water and different doses of saccharin were tested. Saccharin doses were as followed: 0.009%, 0.018%, 0.055%, 0.09%, 0.18%, 0.36%, 0.54%, and 0.9% and this was compared to the consumption of water. During this procedure (Fig. 3), all rats developed a stronger consumption of saccharin as doses increased (see. 0.055% vs. 0.18%) (F_2,19_ = 18.98, p < 0.001) (Fig. 3A) and a decreased consumption as saccharin became aversive (see. 0.18% vs. 0.9%) (F_2, 19_ = 24.89, p < 0.01) showing the specificity of saccharin consumption due to its palatability. PD demonstrated a higher consumption through the levels of saccharin. Indeed, they showed a higher consumption than FAST GD for the doses of 0.055%, 0.09%, and 0.54% (F_2,19_ = 23.64, p < 0.05) and from all rats at the doses of 0.18% and 0.36% (F_2, 19_ = 19.73, p < 0.01). This saccharin consumption was much higher than the consumption of water, for all rats, stable throughout the daily sessions (F_2, 19_ = 28.62, p < 0.001). The basal water consumption was stable before (F_2, 19_ = 5.87, ns) and after (F_2, 19_ = 3.22, ns) the dose-curve effect of saccharin (Fig. 3B). As a result, all rats developed a strong preference for saccharin as doses increased (see. 0.055% vs. 0.18%) (F_2, 19_ = 27.98, p < 0.01) (Fig. 3C).

**Figure 3:**
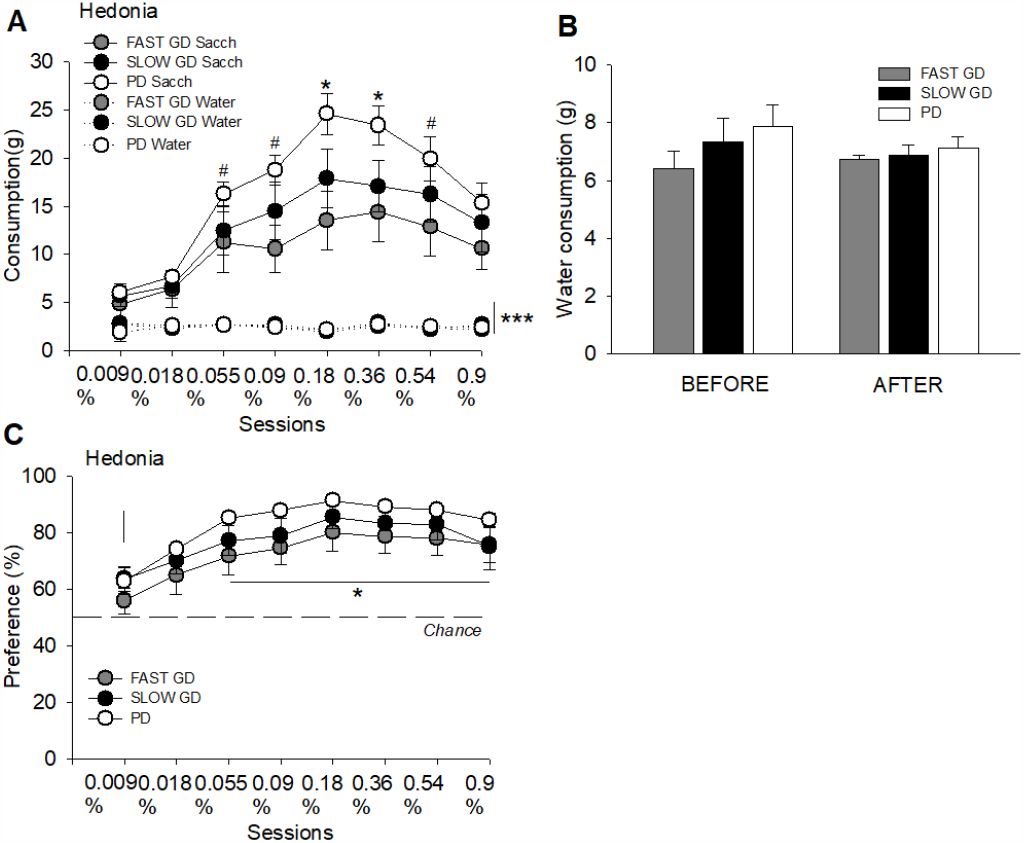
Saccharin intake level (hedonia). (A) Saccharin dose-curve effect for FAST and SLOW GD, and PD. PD consumed more saccharin from the 0.055% dose to 0.54% dose. (B) Water consumption was similar between all groups, and (C) Saccharin preference during the dose-curve experiment. All rats developed a saccharin preference from the 0.054% dose. *\* p < 0*.*05 (comparison all rats), # p < 0*.*05 (comparison FAST GD and poor PD). ANOVA*.

### Inter-individual differences in FR5 and PR2, food-deprived or ad libitum

During this procedure of FR5 (Fig. 4A), rats were not different at the min dose or the max dose, when food-deprived (dose min: F_2,19_ = 5.98, ns; dose max: F_2, 19_ = 4.76, ns). However, when *ad libitum*, a strong difference appeared, PD showed a higher number of lever presses than GD, at the min (F_2, 19_ = 28.78, p < 0.05) and max (F_2, 19_ = 24.90, p < 0.01) doses. Surprisingly, no difference between GD and PD during the PR2 (Fig. 4B) was detected whether animals were food-deprived or *ad libitum*, at the min (food-deprived: F_2, 19_ = 6.56, ns; ad libitum: F_2, 19_ = 4.98, ns) and max (food-deprived: F_2, 19_ = 5.67, ns; ad libitum: F_2, 19_ = 9.97, ns) doses. However, the saccharin consumption at the max dose was lower when all rats were *ad libitum* as compared to the food-deprived condition (F_2, 19_ = 26.76, p < 0.05).

**Figure 4:**
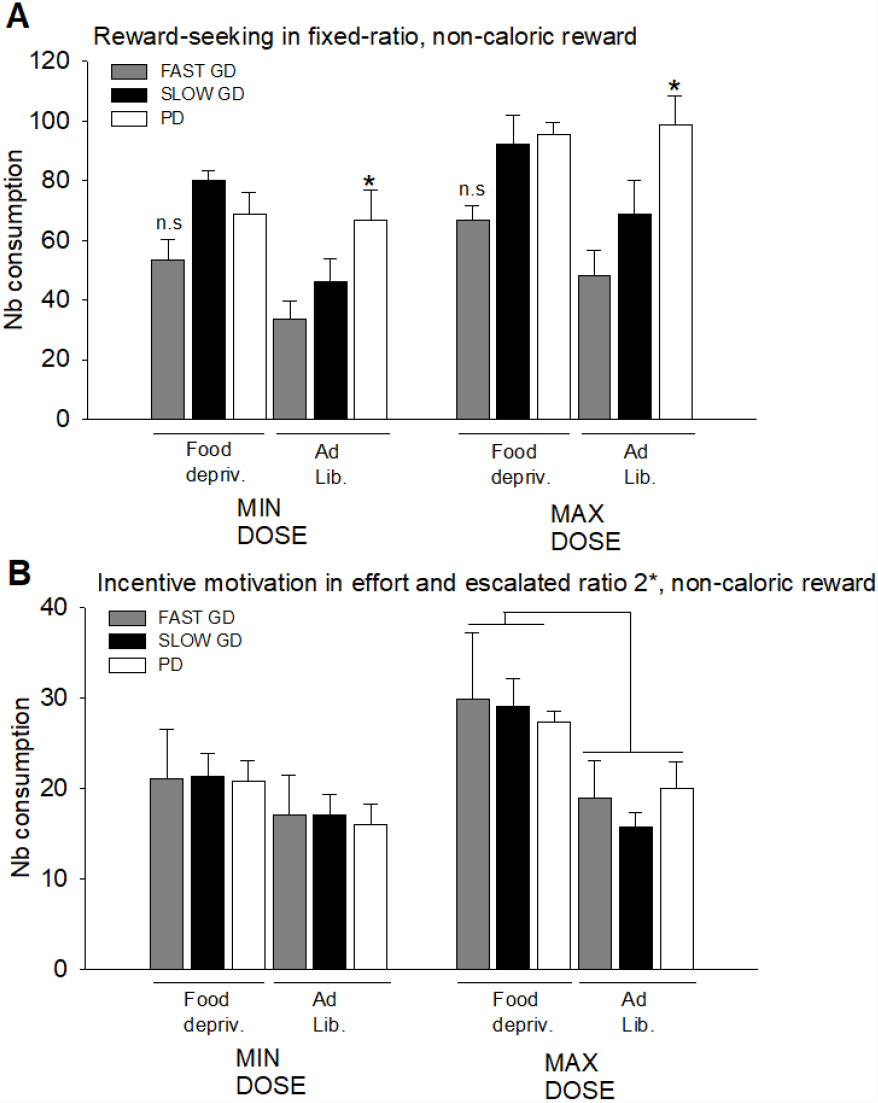
Reward-seeking and incentive motivation evaluation. (A) PD displayed a higher reward-seeking behavior in FR5 at the minimum and maximum doses *ad libitum* while (B) incentive motivation was not affected during PR2 whether food-deprived or *ad libitum. * p < 0*.*05, ANOVA*.

Analyzing qualitatively and separately variables of interest (Fig. 5), certain considered parameters were correlated. In fact, consummatory behavior (hedonia) at the optimal dose (Fig. 5A) was correlated with the 10 first choices (DM, exploration) (r = 0.40, p = 0.067). However, the reward-seeking measure (Fig. 5B) did not correlate with DM (exploration) (r = 0.48, p = 0.05, ns) indicating that the functional link between hedonia and DM is more robust as compared to the aforementioned analyzed correlative measurements. But expectedly, and to continue our assumption, consummatory behavior at the optimal dose (Fig. 5C) was correlated with the reward-seeking measure (FR5 at the optimal dose, *ad libitum*) (r = 0.5, p < 0.05).

**Figure 5:**
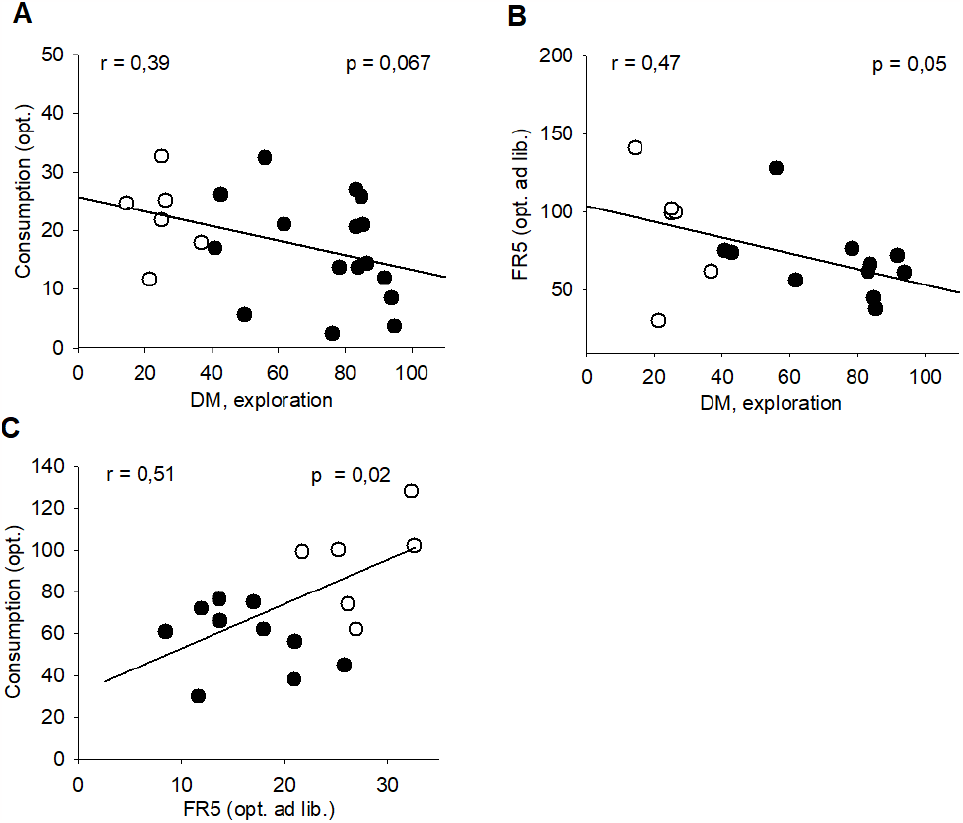
Reward and decision-making correlations. (A) Negative pairwise correlation between consummatory behavior and decision-making (DM, exploration, 10 first choices), (B) No correlation between reward-seeking behavior (FR5) and DM (exploration, 10 first choices), and (C) Positive pairwise correlation between consummatory behavior and reward-seeking behavior (FR5). White circles refer to PD and black circles to PERF. *Pearson correlation (r)*.

The main interest of this study was to decipher the relationship between DM and working memory, aside from the behavioral functions involved in poor DM. We found no correlated measures between WM parameters and DM (exploration) as well as DM (exploitation) (Fig. 6A, B and C) (r = 0, 06, p = 0,8, ns; r = 0, 26, p = 0, 23, ns) suggesting a non-direct relationship between those if any. It supports a non-direct relationship between working memory performance and DM abilities at the time to explore concurrent options in dedicated conflictual and uncertain situations or when establishing preference from the sampled options.

**Figure 6:**
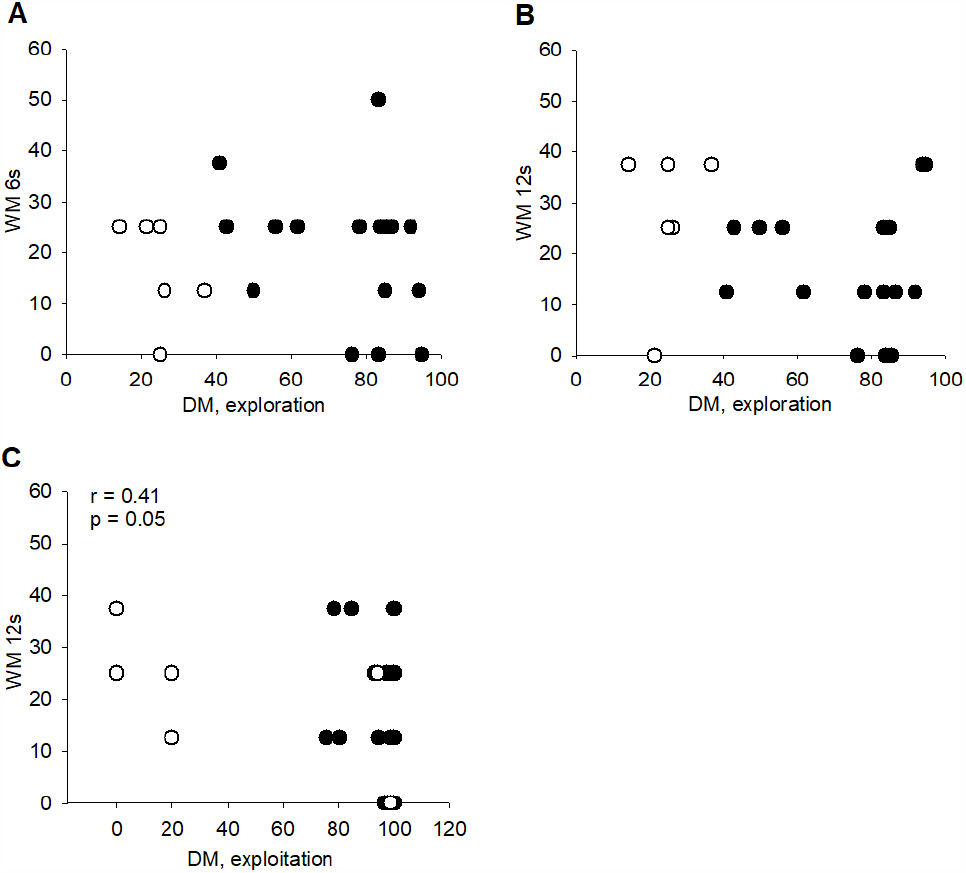
Working-memory and decision-making correlations. (A) No correlation between working memory (WM) (retention delay 6s) and DM (exploration), (B) No correlation between WM (retention delay 12s) and DM (exploration), and (C) No correlation between WM (retention delay 12s) and DM (RGT choices, exploitation). White circles refer to PD and black circles to GD. *Pearson correlation (r)*.

### Inter strain differences, Wistar Han, Lewis and F344

Inter-individual differences in DM as previously shown (Fig. 7). All Wistar Han rats (n = 10), Lewis rats (n = 12) and F344 rats (n = 7) initially selected the various options randomly as reported previously in this experiment before developing a preference for advantageous or disadvantageous options (Fig. 7A, B and C): they clustered into 2 distinct categories depending on their final preference: (1) a majority of GD (Wistar Han rats: GD, n = 6; Lewis rats: GD; n = 6; F344 rats: GD, n = 5) with a strong preference for advantageous options (70 % preference); and (2) a minority of PD (Wistar Han rats: PD, n = 6; Lewis rats: PD; n = 4; F344 rats: PD, n = 2) that persevered in choosing disadvantageous options (70 % preference). Importantly, all rats sampled the 4 options during the first 10 min and experienced the long penalties at least once as reported previously in this experiment or other published work (Fitoussi et al., 2015). Among GD, rats could be distinguished according to the time course of DM. Some rats (Wistar Han rats: SLOW GD, n = 3; Lewis rats: SLOW GD, n = 3; F344: SLOW GD, n = 1) still chose the options randomly during the first 20 min (comparison with chance level (50 %), t Student = 1.87, ns), whereas others (Wistar Han rats: FAST GD, n = 3; Lewis rats: FAST GD, n = 3; F344: FAST GD, n = 4) promptly orientated their choices toward advantageous choices.

**Figure 7:**
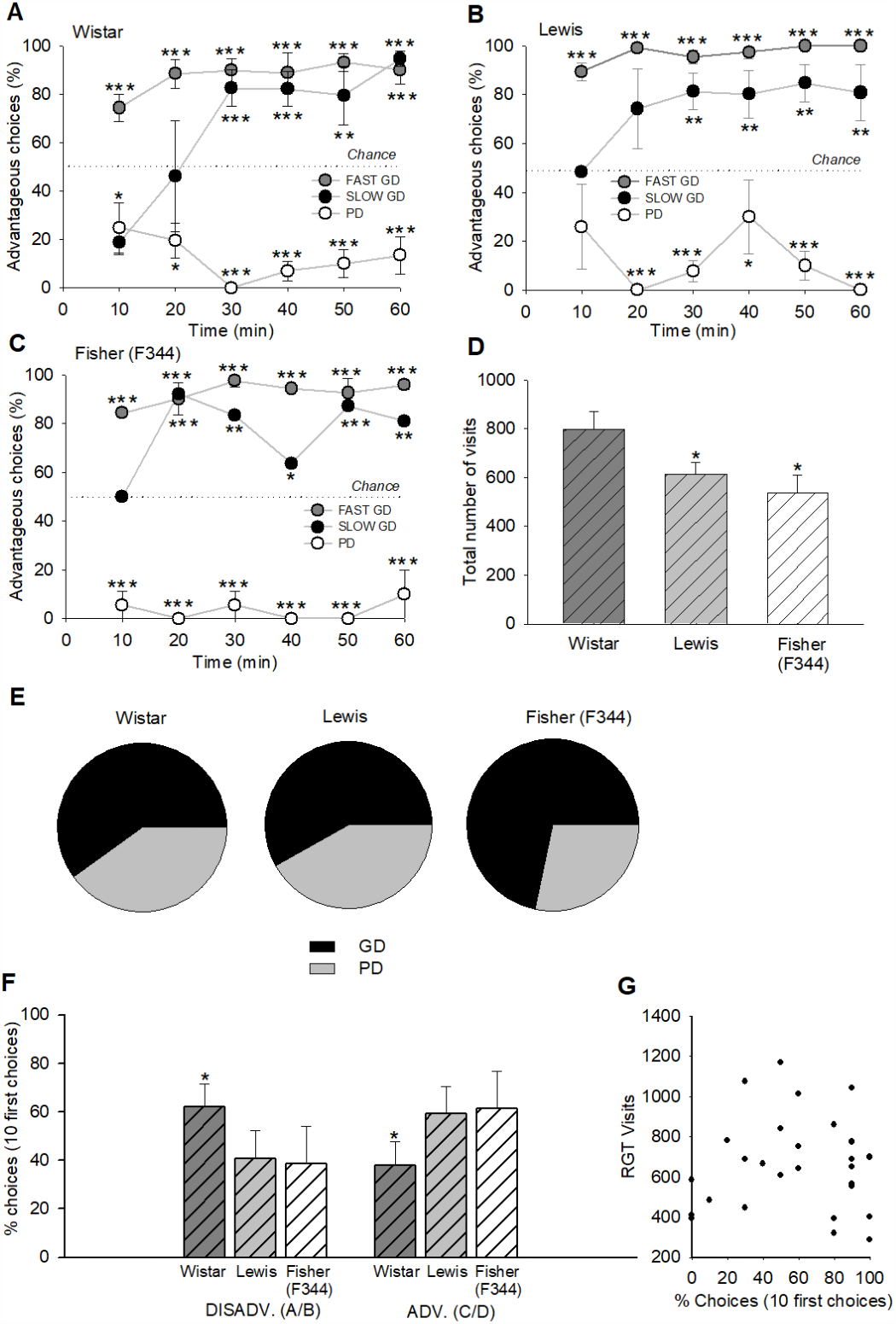
Inbred strain differences in the RGT, Wistar Han, Lewis, and Fischer 344 rats. (A) Pattern of choices of Wistar Han rats, (B) Pattern of choices of Lewis rats, (C) Pattern of choices of Fischer 344 (F344) rats including FAST and SLOW GD, and PD, (D) Total number of visits for the 3 rat strains: Wistar Han rats did a higher number of visits in the RGT, (E) Proportion of GD and PD among the strains: F344 rats displayed a lower number of PD as compared to Lewis rats, and (F) 10 first choice performance for the 3 strains (DM, exploration): Wistar Han rats did poorer performance in the RGT during this early phase, and (G) No correlation between the total number of visits and the 10 first choices (DM, exploration). *\* p < 0*.*05, ** p < 0*.*01, *** p < O*.*001, t-test Student or ANOVA*.

Importantly, strain differences in the total number of visits were reported (Fig. 7D). Indeed, Lewis and Fisher (F344) rats displayed less visits than the Wistar Han rats (F_2,31_ = 27.89, p < 0.05).

The main result of the subpart of this work was that Fisher (F344) rats revealed more GD than Lewis rats. Accordingly, they revealed less PD than Lewis rats (F_2, 21_ = 28.43, p < 0.05). In terms of percentage (Fig. 7E), it appeared that F344 rats demonstrated 71% of GD against 29% of PD, whereas Lewis rats demonstrated 58% of GD against 42% of PD. This is supported by the percentage of choices (10 first choices, i.e., exploration), and the DM variable(s) (Fig. 8F). Indeed, Wistar Han rats poorly performed as compared to Lewis and Fisher (F344) rats (F_2, 21_ = 17.45, p < 0.05) by choosing more A/B options. Surprisingly, RGT (total number of) visits did not correlate with the latter variable (percentage of choices, 10 first choices, exploration) (r = 0.04, ns).

## Discussion

In this study, we aimed at investigating the motivational functions involved in PD as measured in a RGT task (de Visser et al., 2011), as well as the relationship between working memory and DM in this framework. Second, we were investigating the repartition of PD in two inbred strains of rats showing substantial modifications in the dopaminergic system.

Because PD individuals in the RGT (healthy population) displayed higher motivation in PR only when two pellets were delivered in each trial (Rivalan et al., 2009), we sought to investigate the motivational functions in these rats. We reported a dissociation between saccharin intake/hedonia, reward-seeking behavior in FR5, and incentive motivation in PR2 when the reward was saccharin, a non-caloric reward. It is noteworthy that this dissociation was reported in the literature but not for saccharin which represents a good index for vulnerability to drug abuse (Caroll et al., 2008; Holtz et al., 2015). In addicts, it is well-known that *wanting* or *desire* for drugs increases with time, but not hedonia (Barbano and Cador, 2006; Berridge et al., 2004). However, one study reported a linear relationship between sucrose intake, a caloric reward, and incentive motivation in PR (Brennan et al., 2001). PD as measured in the RGT are sensitive to the deprivation state as reward-seeking behavior did not increase when food-deprived as compared to the *ad libitum* condition. This internal state (food-deprived or *ad libitum*) was not enough to drive incentive motivation in PR. Overall, the hedonic set-point (Berridge and Kringelbach, 2010) of PD is higher for a non-caloric reward that does not drive incentive motivation in such conditions. Integrating these notions to the RGT, it favors a reward magnitude variable that drives the motivation of PD, and no metabolic bias that accounts for such motivation. This is also supported by previous work demonstrating the lack of modified performance in the RGT when the food deprivation rate was manipulated (Rivalan et al., 2009). A subtle cost/benefit balance partially drives poor decisions in the RGT, in healthy individuals (Swithers et al., 2010).

No direct relationship between working memory and DM capacities was reported in this experiment. Although surprising in animals, it supports human findings. The implication of working memory (WM) to real-life DM as measured in the Iowa Gambling Task (IGT) has been debated in the literature (Brand et al., 2006). Indeed, some studies revealed that a low WM load could be associated with IGT performance, while others reported no obvious relationship (Bagneux et al., 2013; Brand et al., 2006). Most researchers assumed the existence of an asymmetric relationship between WM and DM, that is, impaired IGT performance does not obligatory rely on WM, but impaired WM leads to poor IGT performance (Bagneux et al., 2013). This was also supported by the effects of dorsolateral prefrontal cortex lesions, a human brain region critically involved in WM (Li et al., 2021; Petrides, 2000, Funahashi, 2006; Goldman-Rakic, 1996; Fuster and Alexander, 1971), and imaging studies (Ernst et al., 2002; Li et al., 2010). It suggests the existence of distinct decision mechanisms within the prefrontal cortex as well as executive functioning (Bechara and Martin, 2004). In animals, it has been shown that intelligence and learning abilities covary with working memory (Dudchenko, 2004; Matzel and Kolata, 2010; Kolata et al., 2005), as the IGT and RGT involved a temporary phase of learning (Fellows 2004; Bagneux et al., 2013). Our work favors an asymmetric relationship, but no working memory impairment associated with poor RGT performance in healthy individuals. It is noteworthy that additional investigations may shed the light on this question in other WM paradigms (Dudchenko, 2004). In the operant chamber configuration, a dissociation may arise as a trend was observed in the 8-Radial Maze, and perseverative behavior was quantified that confirmed previous data collected in Wistar Han rats. Especially, they displayed perseverative-like responding and motor impulsivity (Rivalan et al., 2013).

The second goal of this study was to determine the proportion of PD in two inbred strains of rats, Lewis and F344, that have been proposed as a genetic model for the vulnerability to drug addiction. As it was previously demonstrated that Wistar Han rats displayed a combination of behavioral traits reminiscent of addiction (Rivalan et al., 2013; Rivalan et al., 2009), it was relevant to examine this population in these strains of rats. Moreover, it was shown that a critical set of genes could be involved in addiction (not only one gene), and 40–60% were involved in the variation to the responsiveness of drugs of abuse (Cadoni, 2016). As such, these inbred strains of rats are critical candidates for the vulnerability to addiction, and in our experimental context, to DM evaluation. No study had investigated this relationship. Since we found that there was less PD in F344 rats as compared to Lewis rats, it implies that some neurobiological markers could be involved in DM capacities in the RGT, especially dopamine (Cadoni, 2016). It has been shown that Lewis rats are more sensitive to the reinforcing effects of drugs, using conditioned place preference (Kosten et al., 1994; Guitart et., 1992), operant self-administration (Martin et al., 1999; Ambrosio et al., 1995), and intracranial self-stimulation paradigms (Lepore et al., 1996, Chen et al., 1991). Additionally, these inbred strains of rats displayed substantial variation in the dopaminergic transmission (mesolimbic), essentially via *in vitro* studies (Haile et al., 2001; Harris and Nestler, 1996; Beitner-Johnson et al., 1991). The nucleus accumbens seems to be a pivotal region where TH and DAT levels play a critical role (Gulley et al., 2007; Flores et al., 1998; Harris and Nestler, 1996). Further, dopaminergic neurons in the ventral tegmental area demonstrated more burst events in Lewis rats as compared to F344 rats. It supports our previous work that reported higher dopaminergic levels at rest in (healthy) PD (Fitoussi et al., 2015) although the functionality of the nucleus accumbens of Lewis and F344 rats is not entirely known.

One important result of this study is the improved performance of Lewis and F344 rats early in the task as compared to Wistar Han rats. It has been shown that this critical phase corresponds to the exploration, a dedicated step for sampling the available options (Berntson et al., 2011; Craig et al., 2009). It also involved a part of learning and attractiveness for food pellets. The fact that two inbred strains of rats are better performers than Wistar Han rats is presently unknown. Usually, Lewis rats displayed higher striatal dopamine levels as compared to F344 whereas Wistar Han rats would exhibit an intermediate level. It is noteworthy that dopaminergic levels in prefrontal regions, especially the infralimbic cortex could explain, at least in part, this phenomenon (Fitoussi et al., 2015). Additionally, this improved performance parallels the lower number of visits (as compared to Wistar Han rats) and pointed out a dopaminergic component (Carker et al., 2013), striatal or prefrontal.

It is noteworthy that Lewis and F344 rats exhibit some significant differences in anxiety and stress behaviors. Indeed, it was shown that Lewis rats could exhibit a higher anxiety level than F344 (Cadoni, 2016). Because Wistar Han rats poor decision-makers display significant differences in risk-taking behavior as measured in the Elevated-Plus-Maze (Rivalan et al., 2009) but not in anxiety level, we believe that the most significant differences observed in this study was not related to any variations in anxiety level and stress, but rather, variations in PD repartition among these strains was related to a predominant dopaminergic component. Moreover, differences in anxiety among Lewis and F344 rats were not systematically reported, thus it leads to contrasted results in these strains. Overall, these results supported the current literature and expected data regarding DM capacities in these inbred strains of rats.

In summary, this work completes the preclinical panel of behavioral functions that relates to poor DM, and extends a presumed role of dopamine in such processes.

## Acknowledgments

I would like to thank Dr. Françoise Dellu-Hagedorn and Dr. Martine Cador for their conceptual help on this work. This work was supported by the CNRS (French National Research Council).

